# Top-Down Modulation of Visual Action Perception: Distinct Task Effects in the Action Observation Network

**DOI:** 10.1101/2024.06.14.598806

**Authors:** Aslı Eroğlu, Burcu A. Urgen

## Abstract

Perceiving others’ actions is essential for survival and social interaction. Cognitive neuroscience research has identified a network of brain regions crucial to visual action perception, known as the Action Observation Network (AON), comprising the posterior superior temporal cortex (pSTS), posterior parietal cortex, and premotor cortex. Recent research highlights the importance of integrating top-down processes, such as attention, to gain a deeper understanding of action perception. This study investigates how attention modulates the AON during human action perception. We conducted a two-session fMRI experiment with 27 participants. They viewed eight videos of pushing actions, varying in actor (female vs. male), effector (hand vs. foot), and target (human vs. object). In the first session, participants focused on specific features of the videos (actor, effector, or target). In the second, they passively viewed the videos. From the passive viewing session data, we defined regions of interest (ROIs) in the pSTS, parietal, and premotor cortices for each hemisphere. We then performed model-based representational similarity analysis (RSA) and decoding analysis. RSA results showed that only the task model, among all tested models, exhibited a significant correlation with neural representational similarity matrices (RDMs) across all ROIs, indicating a specific alignment between AON nodes and the ongoing task. Decoding analysis further showed that different task types uniquely affected each AON node, indicating feature- and region-specific interactions. These findings underscore that top-down attentional processes significantly alter neural representations within the AON, highlighting the dynamic interplay between attention and action perception in the brain.

## 1. Introduction

Perception of others’ actions is one of the core abilities to navigate and comprehend the external world. Its evolutionary roots extend deep into the animal kingdom, which serves as a vital survival tool, allowing organisms to identify opportunities and threats in their environment. For humans, action perception goes beyond mere survival; it also plays a central role in our social interactions. Our capacity to interpret the actions of others has been the cornerstone of effective communication, collaboration, and self-preservation.

Years of cognitive neuroscience research show that visual perception of actions is supported by a network of regions including the early visual cortex and motion-sensitive regions like MT+, extrastriate body area (EBA) (Grossman & Blake, 2002, Peelen et al., 2006, Jastorff & Orban, 2009) which then extend into the Action Observation Network (AON) (Saygin, 2007; Caspers et al., 2010; Nelissen et al., 2011). This network’s core regions include the posterior superior temporal cortex (pSTS), the posterior parietal cortex, and the premotor cortex (Figure 1).

**Figure 1:**
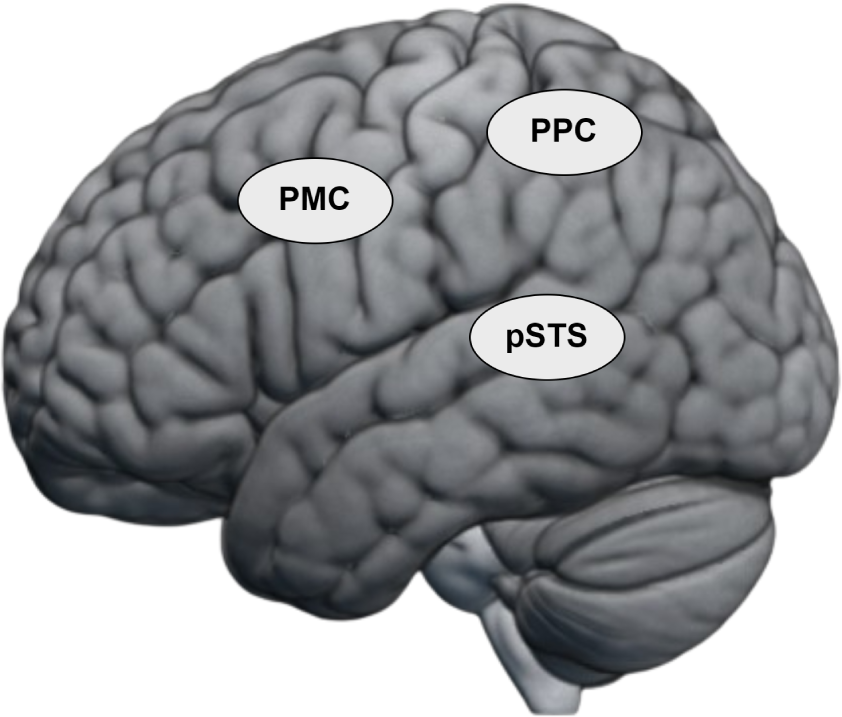
The core regions of the Action Observation Network: posterior superior temporal cortex (pSTS), posterior parietal cortex (PPC), premotor cortex (PMC)

The pSTS is one of the brain regions that consistently shows activity in neuroimaging studies of action observation. Given its anatomical proximity to the early visual cortex (EVC), this area is usually considered to serve as the initial site for visual information flow into the AON. Several studies suggest that pSTS integrates the motion information conveyed through the dorsal pathway and the shape information through the ventral pathway (Vaina et al., 2001, Giese & Poggio, 2003). In addition, lesion mapping studies in humans have indicated a higher likelihood of difficulties in action perception in individuals with damage specifically in the STS region compared to other areas (Saygin, 2007). It is well-established that the pSTS region demonstrates activation specifically when processing social stimuli (e.g., faces, biological motion) in contrast to other stimuli (e.g., objects, scrambled motion) (Watson et al., 2014; Saygin et al 2004). It also shows greater activation when humans perform biological motion compared to animals (Kaiser et al., 2012). Furthermore, the pSTS region is sensitive to complex social cues such as gaze direction (Senju & Johnson, 2009) and head orientation (Carlin & Calder, 2013). Thus, pSTS emerges as a vital hub, not just for integrating visual details in action perception but also for decoding complex social cues (Deen et al, 2015; Wurm et al, 2017; McMahon & Isik, 2024), emphasizing its dual role in understanding both action observation and social cognition in the brain. Based on these socially important functional properties, it has been indicated as a core region of the recently proposed third visual pathway of the visual system (Pitcher & Ungerleider, 2021).

The posterior parietal cortex (PPC) is another crucial region within the AON, responsible for encoding higher-level visual and abstract information related to actions and forming a coherent action model. The PPC utilizes visual representations conveyed from pSTS and encodes interactions between the target of action and the involved body parts during execution (Urgen et al., 2019; Jastorff et al., 2010). Additionally, the PPC encodes information about the purpose and context of actions, such as the goal or intention behind the action (Gallivan et al., 2011; Urgen et al, 2019). Individuals with parietal damage may encounter difficulties not in perceiving actions visually, but in grasping the meaning and appropriateness of actions within a given context (Binder et al., 2017). This emphasizes the pivotal role of the parietal region in shaping abstract action representations. The PPC’s capacity to encode information about the reasons and contextual aspects of an action bridges the visual and abstract components essential for constructing the meaning of an action. Consequently, it contributes to a comprehensive understanding of not only the physical movements involved in an action but also the underlying intention, goal, and contextual relevance of the action.

The premotor cortex, located within the frontal cortex, is the third important region in the AON. This region hierarchically lies above the pSTS and parietal regions within the AON (Nelissen et al., 2011). The premotor cortex, as revealed by neuroimaging studies, demonstrates a specialized sensitivity to body parts involved in action execution (Di Dio et al., 2013, Fabbri et al., 2016). This sensitivity aligns with its well-established functional organization, where its distinct sections are dedicated to controlling specific body parts, underscoring its pivotal role in coordinating a diverse range of motor actions (Fujii et al., 2008; Jastorff et al., 2010). The premotor cortex exhibits a somatotopic organization, reflecting the representation of different body parts. Notably, the dorsal premotor cortex is responsible for representing movements involving more distant body parts, such as finger movements, while ventral regions encode movements of closer body parts, like shoulder movements (Sakreida et al., 2005). Furthermore, some studies provide evidence supporting the involvement of the premotor cortex in the planning and execution of actions (Krams et al., 1998; Binkofski et al., 1999). In summary, these findings collectively emphasize the crucial role of the premotor cortex within the AON, where it participates in the planning, coordination, and execution of actions.

Neuroimaging studies often explore the AON by passively observing actions, but this method overlooks the complex interplay between bottom-up and top-down processes crucial for real-life perception (Kay et al., 2023). Recent research on action observation emphasizes integrating top-down processes like expectation, prediction, and attention, which significantly shape perception (Kilner et al., 2007; Urgen & Miller, 2015; Urgen & Saygin, 2020). Attention, in particular, which is crucial for directing cognitive resources efficiently, influences various cognitive processes and can enhance neural responses to attended stimuli, thus impacting behavioral performance (Borji and Itti, 2012; Cohen & Maunsell, 2009). In particular, feature-based attention modulates neural activity in favor of attended features and suppressing unattending features, revealing the complexity of attentional mechanisms in perception (Maunsell & Treue, 2006; Treue, 2003). Incorporating top-down attention tasks in action observation studies is critical for a comprehensive understanding of visual perception of actions, especially within real-life contexts.

There are only a handful of studies investigating how the attention mechanisms modulate the AON. One such study conducted by Stehr et al. (2021) investigated how action coding in the pSTS region changed under different attention conditions using functional magnetic resonance imaging (fMRI). Their findings revealed that directing attention to action categories significantly enhanced the distinctiveness and categorization of neural responses within the right pSTS. This suggests that attention not only influences the overall activity in the pSTS but also shapes the way neural populations represent different actions. These findings highlight the significance of top-down processes in shaping the perception.

In an fMRI study conducted by Orban and colleagues (2019), participants observed a set of videos in which an actor manipulated an object while engaging in tasks that involved distinguishing between action categories (pushing or pulling the object), identifying the object’s color, or recognizing the performing actor’s identity. Within the Region of Interest (ROI) analyses, researchers included certain regions such as the lateral-occipital regions near the pSTS, the parietal cortex, and the premotor cortex. The results suggest that participants’ brain activities varied depending on the attention tasks, even when viewing the same videos. Specifically, when participants engaged in the action discrimination task, activation was observed in the parietal cortex, contrasting with the lack of activation in other discrimination tasks. Additionally, the authors stated that lateral-occipital areas showed activation in a non-task-dependent manner, while the premotor region exhibited task specificity. This suggests that not every attention task equally influences every brain region; rather, different tasks affect brain regions to varying degrees.

Another significant study supporting the idea that top-down attention processes can influence action perception was conducted by Shahdloo et al. (2022). This study addresses the brain’s representation of diverse visual action categories and examines how these representations dynamically change depending on the task performed. Participants watched a series of videos containing numerous visual action categories (e.g., ‘hitting’ or ‘lifting’). They performed a task requiring a covert search for the target action categories, while their brain activity was measured using fMRI. In summary, the article demonstrates that a natural visual search for specific action categories influences semantic representations in the brain, causing tuning shifts in neural activity toward the target category, within and beyond the AON. This suggests that dynamic attentional mechanisms enhance action perception by effectively allocating neural resources to emphasize task-relevant action categories, providing insights into how humans perceive others’ actions in dynamic daily life experiences.

As evident from the examination of the limited studies conducted in this field, research on the impact of top-down attention processes on action perception has generally focused on manipulated objects, actors, and action classes. However, it is known that the body parts used during the execution of an action also play a significant role in constructing the meaning of the action. This feature is processed in the premotor cortex and the parietal cortex of the AON (Jastorff et al, 2010), and directed attention to the effector can specifically influence these regions. Furthermore, the target to which the action is directed is another crucial aspect affecting the meaning of the action. For example, when the target is an object, a pushing action is considered a manipulation action, while if the target is a human, it becomes a social action with a certain valence value. Despite their relevance, the impact of attention tasks on the perception of specific action features (e.g., actor, effector, target) remains unexplored in the action perception literature. Additionally, no research has systematically manipulated attention to these features to understand their interplay.

This study aims to investigate the dynamic interaction between attentional processes and the AON across the pSTS, parietal cortex, and premotor cortex. Specifically, we seek to understand how attention modulates these regions during the perception of human actions, building upon prior neuroimaging studies. To achieve this, our methodology involves functionally localizing core AON regions using univariate analysis in a passive viewing session where participants observed the action videos. Subsequently, we apply univariate, multivariate pattern analysis (MVPA), and representational similarity analysis (RSA) on the data from active sessions, where participants engage in attention tasks. We aim to investigate whether attention directed towards the actor, effector, or target features of an action differentially modulates activity across the AON. This research may reshape our understanding of action perception, suggesting a more complex interplay between attention and neural processing than previously assumed.

## 2. Methods

### 2.1 Participants

30 healthy individuals participated in the two-session fMRI experiment. They had either normal or corrected-to-normal vision and were not taking medication for neurological or psychiatric conditions. 3 of the participants were excluded due to excessive motion and technical problems. The remaining 27 participants included 12 females with the age range of 21-31 and a mean age of 24.5. The study was approved by the Human Research Ethics Committee of Bilkent University and each participant provided written informed consent and filled the MRI prescreening form.

### 2.2. Stimuli, Experimental Design and Procedure

#### 2.2.1 Stimuli

Eight action videos each lasting 3 seconds were recorded. All videos depicted a “pushing” action but varied in terms of three parameters: the actor performing the movement, the body part (effector) used for the action, and the target of the action. The actions were performed by two different actors, one female and one male. Each actor performed the pushing action using two different effectors, hand and feet. Finally, the actors directed the pushing action toward two different targets, a human and an object.

The videos were recorded with the Sony HDR-CX405 Handycam Video Camera at 50fps and 1280×720 resolution. All videos were recorded from the same distance and the actors’ positions were kept fixed and symmetrical. The actors were instructed to minimize any variation in their movements and maintain neutral facial expression and they practiced the movements before the filming took place/videos were recorded.The clothing of the two actors was chosen to be similar to each other (black top, black pants).

#### 2.2.2 Experimental Design and Setup

The study consisted of two fMRI sessions. The first session was an ‘active’ session where participants watched the video stimuli under task conditions and did evaluations about the video. The second session was a ‘passive’ session where the same stimuli were presented without any task.

The active session was composed of 8 runs and each run lasted approximately 8 minutes. Each fMRI run included three distinct attention tasks—**Actor, Effector**, and **Target**—each referred to as a task block (Figure 2). Within each run, each block type was presented twice, resulting in a total of 6 block presentations. Three attention blocks were grouped together to form one set, and each set was shown twice, with the order of blocks randomized within each set.

**Figure 2.**
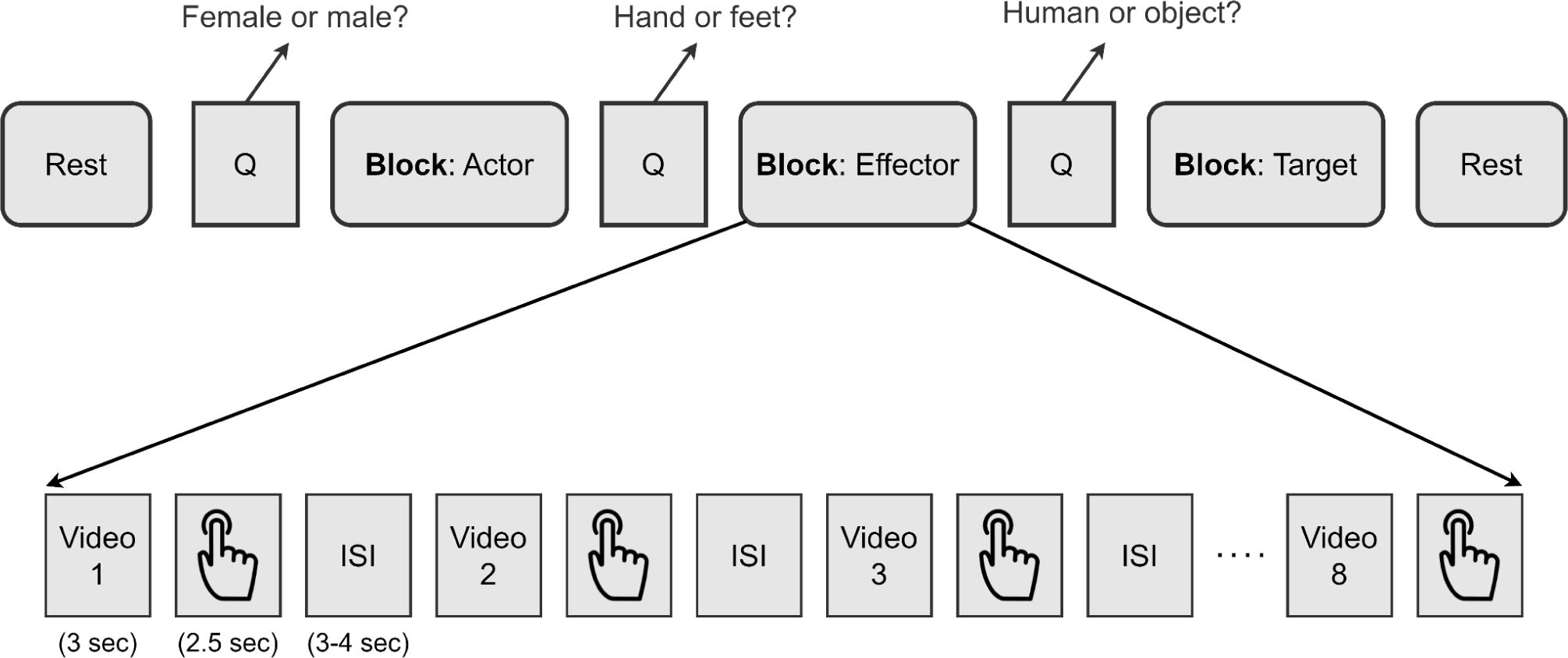
Experimental design for the active session, showing one set (top) and one block (bottom). ISI: Interstimulus interval.

Before each block, participants were presented with an instruction screen (4s) indicating the task they would perform in that block (Figure 2, top section, ‘Question’ screen). Based on the question presented, participants evaluated the videos during the block using a binary button after they watched the video. In the Effector block, they evaluated whether the action was performed with the hand or the feet. In the Actor block, they evaluated whether the actor was female or male, and in the Target block, they evaluated whether the action was directed towards a human or an object. Before and after each set in each run, there was a 12-second rest block.

Within each block, 8 different video stimuli were randomly presented (Figure 2, bottom section). After each video, participants were given 2.5 seconds to answer the question. Following this, interstimulus intervals (ISI) between 3 and 4 seconds were shown, during which participants fixated on the fixation cross. The average duration of the ISIs within each block was 3.5 seconds.

A practice session was conducted to ensure participants’ understanding of the experiment. In this mini-experiment, each block was shown once. Instead of displaying all eight videos, two videos were carefully selected to be shown in the blocks. These two chosen videos cover all the different variations (one video consisted of the “female, hand, and object” condition, while the other is from the “male, feet, and human” condition).

In the passive session, there were 4 runs each lasting approximately 6 minutes. The participants viewed the same set of stimuli, without any required response or task to perform. In each run, they passively viewed a block of 8 videos, with each block repeated six times. This resulted in each video being presented a total of 24 times. Data from this session was used to examine the effects of attention tasks, as well as to extract ROIs within the AON to perform further analysis.

Video stimuli in the MRI machine were presented to the participants through the TELEMED screen, which they viewed through a mirror mounted on the head coil (∼10cm distance between the mirror and eyes). The TELEMED screen had a refresh rate of 60 Hz and a resolution of 1920×1080. The videos were displayed on the screen in a size of 550×340 pixels, with a uniform gray background. Stimulus presentation and response collection codes were written using the PsychoPy library and custom functions in Python.

### 2.3. MRI Data Acquisition

Participants’ brain activity was recorded using a 32-channel head coil on a 3T Siemens TimTrio MRI scanner at the National Magnetic Resonance Research Center (UMRAM) at Bilkent University. Soft cushions made of materials that minimize head movement were placed under the participants’ heads, around their necks, and under their legs for support. Earplugs were also provided to reduce exposure to the high noise levels in the MRI machine.

Prior to the functional scans, high-resolution T1-weighted structural scans of participants’ brains were acquired for both experiments (TR=2600 ms, TE=2.92 ms, flip angle=12°, FoV read=256 mm, phase FoV=%87.5, 176 slices, and 1×1×1 mm^3^ resolution). Then, for the active session, 168 functional scans and for the passive session 129 functional scans were obtained using gradient echo-planar imaging (TR=3000 ms, TE=22 ms, flip angle=90°, 64 x 64 matrix, FoV read=192 mm, 43 slices with a thickness of 2.5 mm and a resolution of 3×3×2.5 mm^3^).

#### 2.3.1 Anatomical and Functional Data Preprocessing via fMRIPrep

Results included in this manuscript came from preprocessing performed using fMRIPrep 21.0.1 (Esteban et al., 2019), a Nipype (K. Gorgolewski et al., 2011) based tool. Each T1w (T1-weighted) volume was corrected for INU (intensity non-uniformity) using N4BiasFieldCorrection v2.1.0 (K. J. Gorgolewski et al., 2017) and skull-stripped using antsBrainExtraction.sh v2.1.0 (using the OASIS template). Brain surfaces were reconstructed using recon-all from FreeSurfer v6.0.0 (Dale et al., 1999), and the brain mask estimated previously was refined with a custom variation of the method to reconcile ANTs-derived and FreeSurfer-derived segmentations of the cortical gray-matter of Mindboggle (Klein et al., 2017). Spatial normalization to the ICBM 152 Nonlinear Asymmetrical template version 2009c (Fonov et al., 2009) was performed through nonlinear registration with the antsRegistration tool of ANTs v2.1.0 (Avants et al., 2008), using brain-extracted versions of both T1w volume and template. Brain tissue segmentation of cerebrospinal fluid (CSF), white-matter (WM) and gray-matter (GM) was performed on the brain-extracted T1w using fast (Zhang et al., 2001) (FSL v5.0.9).

Functional data was slice time corrected using 3dTshift from AFNI v16.2.07 (Cox, 1996) and motion corrected using mcflirt (FSL v5.0.9 (Jenkinson et al., 2002)). This was followed by co-registration to the corresponding T1w using boundary-based registration (Greve & Fischl, 2009) with 9 degrees of freedom, using bbregister (FreeSurfer v6.0.0). Motion correcting transformations, BOLD-to-T1w transformation and T1w-to-template (MNI) warp were concatenated and applied in a single step using antsApplyTransforms (ANTs v2.1.0) using Lanczos interpolation.

Physiological noise regressors were extracted applying CompCor (Behzadi et al., 2007). Principal components were estimated for the two CompCor variants: temporal (tCompCor) and anatomical (aCompCor). A mask to exclude signal with cortical origin was obtained by eroding the brain mask, ensuring it only contained subcortical structures. Six tCompCor components were then calculated including only the top 5% variable voxels within that subcortical mask. For aCompCor, six components were calculated within the intersection of the subcortical mask and the union of CSF and WM masks calculated in T1w space, after their projection to the native space of each functional run. Frame-wise displacement (Power et al., 2014) was calculated for each functional run using the implementation of Nipype.

Many internal operations of fMRIPrep use Nilearn (Abraham et al., 2014), principally within the BOLD-processing workflow. For more details of the pipeline see http://fmriprep.readthedocs.io/en/latest/workflows.html.

### 2.4 Behavioral Analysis

Behavioral analysis of the active session focused on reaction times and accuracy. To assess potential differences in task difficulty, a one-way repeated measures ANOVA was conducted across the three tasks.

### 2.5 Univariate Analysis and Activation Maps

After preprocessing, we first conducted a univariate analysis using the SPM12 software. The analysis employed the general linear model (GLM) in both active and passive sessions.

In the passive session, the GLM included the following regressors: action movies, rest, instruction, ISI, 6 head movement regressors extracted from preprocessing. Functional scans from the passive session were smoothed with a full width at half maximum (FWHM) = 4mm kernel. Then, we contrasted the action movies condition with the rest condition. The resulting contrast map was used to establish the ROIs for further analysis. Six distinct ROIs in the AON, which are pSTS, parietal, and premotor regions in both hemispheres were extracted.

In the active session’s univariate analysis, each task condition was defined as a distinct variable. These variables included 3 attention task blocks, rest, instruction, ISI, response duration, and head movement constants for each run. For further analysis in the active session, two more GLMs were run based on the analysis methods that will be conducted. For the RSA analysis, each video within each task was defined as a distinct variable. For decoding analysis, each trial was treated as a separate variable.

Following the estimation of the beta values in GLM, contrast analysis was performed to compare activation differences among conditions. For the active session, we contrasted each attention block with the rest block. Also, to examine the overall task effect and enable a comparison with the contrast image from the passive session, we contrasted the ‘all tasks’ condition with the ‘rest’ condition. The resulting contrast images for activation maps are smoothed with a 4mm full width at half maximum (FWHM) kernel for visualization purposes.

### 2.6 Representational Similarity Analysis

To explore the representational content within the core nodes of the AON (pSTS, parietal, and premotor cortex), we conducted a model-based representational similarity analysis. The analysis was carried out using the RSA Toolbox (Nili et al., 2014), with some modifications made to the built-in scripts. Contrast images were generated for all 24 regressors (3 tasks x 8 videos) in comparison to the rest. The thresholded SPM images (spmT’s) resulting from this analysis were employed as inputs for the RSA.

To examine the correlations between model and neural representational dissimilarity matrices (RDMs), we generated six distinct models. The first three models covered the actor, effector, and target features of the videos. These models were binary matrices, with a value of 1 indicating dissimilarity and 0 indicating similarity. In the Actor model, the representation of same-gender pairs (female-female and male-male) is encoded as 0, while a mixed-gender pair (female-male) is represented as 1. In the Effector model, same-limb pairs (hand-hand and feet-feet) are denoted as 0, and a mixed-limb pair (hand-feet) is denoted as 1. Finally, in the Target model, identical features, such as object-object and human-human pairs, are represented as 0, while different features are represented as 1. The fourth model was the task model, characterized by again a binary RDM. In this case, identical task values (Actor-Actor, Effector-Effector, and Target-Target) correspond to 0, while different task values (Actor-Effector, Actor-Target, Effector-Target) were denoted as 1. The next model, referred to as the visual model, covered the visual similarities of the eight videos. We employed Matlab’s structural similarity function (ssimval) to assess video similarities. This function considers the luminance, contrast, and structure values of the images, and we performed the averaging over all frames for each video. The resulting output values range from 0 to 1, with 1 indicating complete dissimilarity and 0 indicating perfect similarity. Lastly, we introduced a random model to serve as a control model for the analysis. The structure of model RDMs (A) and a visual representation of models (B) can be found in Figure 3.

**Figure 3.**
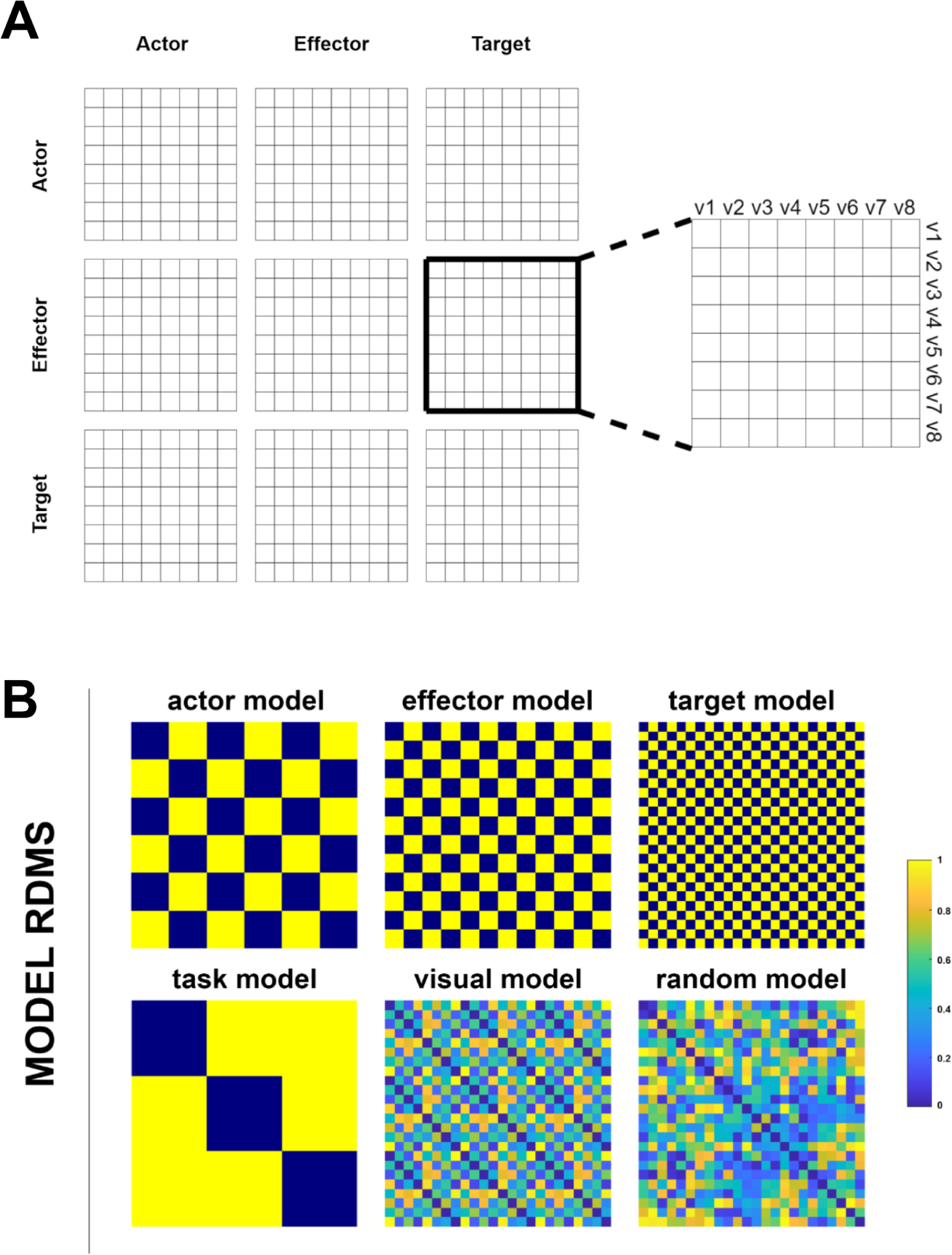
A) The structure of the model RDMs presents a 24×24 matrix, including 8 videos within each of the 3 tasks. **B)** Six model RDMs, to be compared with neural RDMs for each ROI. The actor, effector, and target models represent the features of the videos. The task model includes the attention blocks in the active session. The visual model illustrates how videos are similar based on their low-level visual features. The random model is generated using random numbers between 0 and 1 to serve as a control.

### 2.7 Decoding Analysis

To distinguish neural activity within the ROIs, we employed a decoding analysis using the Decoding Toolbox (Hebart et al., 2015). With this analysis, our primary objective was to identify and characterize distinct features within ROIs under different task conditions. We aimed to classify actor (female vs. male), effector (hand vs. foot), and target (human vs. object) features under each task.

These binary classifications were carried out independently in each hemisphere (left and right), for each ROI (pSTS, parietal, premotor) and across all tasks (actor, effector, target), yielding a total of 54 distinct decoding analyses. The beta images from GLM analysis were used as inputs for our classification algorithms. A linear support vector machine (LibSVM) (Chang & Lin, 2011) was implemented with a fixed cost parameter of c=1. To validate the robustness of our results, a leave-one-run-out cross-validation procedure was employed.

For each classification, decoding accuracy was computed. To assess the success of the decodings, a one-sample t-test was employed, with the hypothesis that the sample mean exceeded 50% for each decoding condition. Then we performed a repeated-measures ANOVA to assess the impact of hemisphere, ROI, task, and features on classification accuracies. This analysis allowed us to examine both main and interaction effects across these factors.

## 3. Results

### 3.1 Behavioral Results

Behavioral results were examined to check whether the participants understood the task and if there was a difference between the tasks in terms of difficulty. The average accuracy rate for all tasks was 95.5% (sd=+-7.9%) across participants. For the actor task, the accuracy was 95.3% (sd =+-7.6%), for the effector task, it was 95.9% (sd =+-8.3%), and for the target task, it was 95.5% (sd=+-8.1%). A one-way ANOVA was performed to compare the accuracy rates of the tasks. The results indicate no statistically significant differences among these tasks (F(2, 78) = 0.04, p = 0.964).

For the response time analysis, the false answers were excluded. The average response time for all tasks was 0.80 seconds (sd=+-0.18 seconds) across participants. For the actor task, the average response time was 0.81s (sd =-+ 0.20), for the effector task, it was 0.78s (sd =+-0.18), and for the target task, it was 0.81 (sd=+-0.17). A one-way ANOVA was performed to compare the response times of the tasks. The results indicate no statistically significant differences among these tasks (F(2, 78) = 0.21, p = 0.812).

### 3.2 Univariate Analysis and Activation Maps

#### 3.2.1 Data Cleaning

Out of the 30 participants who initially participated in the study, one was excluded due to technical problems. Motion regressors were derived from the preprocessing outcome for the remaining 29 participants. The motion exclusion criteria for each run were defined as either having a total movement of one voxel size or half-voxel size spikes. Two participants were fully excluded due to excessive head motion observed during more than half of the total runs. Additionally, for the first session (active session), one run from three participants and two runs from two participants were removed from the analysis. For the second session (passive session), one run from one participant was also excluded from the analysis.

#### 3.2.2 Univariate Results

In the passive session, our videos effectively stimulated the AON, evident from the activation of key regions associated with action observation. At a significance threshold of FWE corrected p<0.05, early visual areas including V1, V2, V3; the regions in the occipito-temporal cortex including the MT cluster and posterior superior temporal sulcus; and the PGp region in the posterior parietal cortex were activated (Figure 4, top). At an uncorrected p<0.001 threshold, additional activations surfaced in the posterior parietal cortex (phAIP, PGp, PGa, PFt, PFpop, PFm, PFcm, PF) (Caspers et al, 2006) as well as in premotor cortex, V7, VIPS, and BA2 region. These regions were illustrated by the white borders in both Figure 4 and Figure 5.

**Figure 4.**
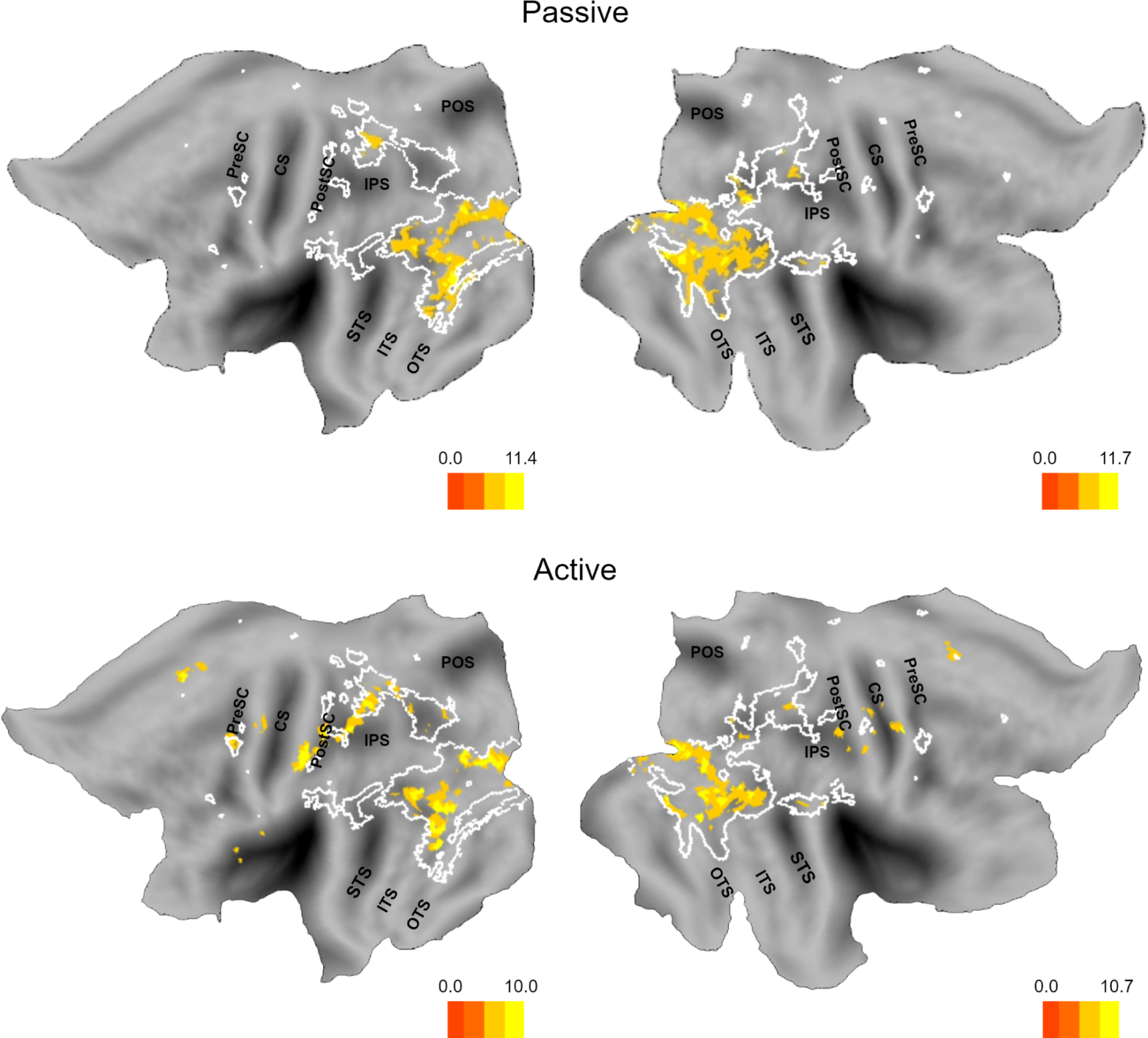
Activation maps of “all videos minus rest” contrast from passive session (top) and active session (bottom). The results are adjusted for significance at p<0.05 with FWE correction. The white borders are from the contrast between the “all videos and the rest” condition from the passive session at p<0.001, uncorrected. Maps are visualized on the flat map using Caret5 software. Landmarks are indicated in black: PreSC (pre-central sulcus), CS (Central sulcus), PostCS (post-central sulcus), IPS (intra-parietal sulcus), POS (parieto-occipital sulcus), STS (superior temporal sulcus), ITS (inferior temporal sulcus), OTS (occipito-temporal sulcus).

**Figure 5.**
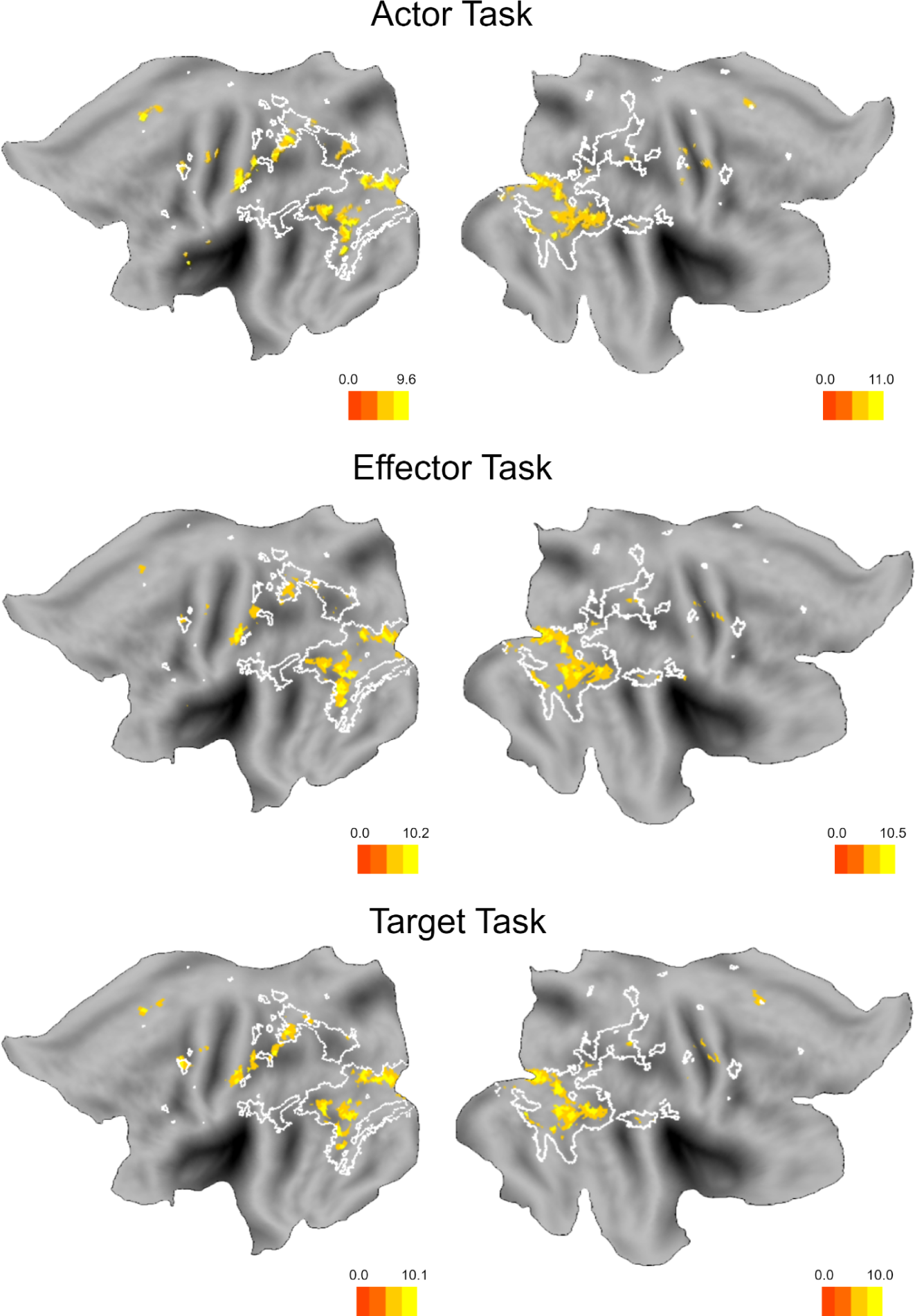
Activation maps of all videos minus rest contrast from the actor task (top), the effector task (middle) and the target task (bottom). The results are adjusted for significance at p<0.05 with FWE correction. The white borders are from the contrast between all videos and the rest condition from the passive session at p<0.001, uncorrected.

In the active session, contrasting all movies against rest (regardless of task) at a significance threshold of FWE corrected p<0.05 yielded similar activation patterns to the passive session, while showcasing extended activation areas, notably within the posterior parietal cortex, premotor, and frontal regions. These supplementary activation regions include DIPSA, DIPSM, phAIP (extended) (Orban, 2016), BA2 (extended), BA3, BA4, PMm, and premotor areas (Figure 4, bottom).

Next, we visualized the activation maps for each task individually. The resulting activation patterns exhibit resemblance to those observed in the active session’s “all tasks minus rest” condition, primarily localizing to regions associated with action observation (Figure 5).

Parietal region activity varies slightly across tasks. Notably, DIPSA is active in all tasks, while DIPSM is active in effector and target tasks within the left hemisphere. In the right hemisphere, DIPSA is active for the actor task. On the other hand, DIPSA and DIPSM show overlapping activity at the border between the two regions for effector and target tasks in the right hemisphere. OP1 region is active only in the effector task within the right hemisphere. In terms of the motor activation area derived from Ferri et. al (2015), the left hemisphere displayed activity during all tasks, while it is inactive in the right hemisphere. Also, in the left hemisphere we observed activity in premotor dorsal (PMd), premotor medial (PMm) and premotor ventral (PMd) regions. In the right hemisphere, premotor activity was less pronounced. We only observed activity in PMm for the actor and target task. In the lateral occipitotemporal cortex, all three tasks activate areas on the middle temporal gyrus (MTG), MT+, and occipito-temporal sulcus (OTS), with very similar activation patterns for both hemispheres.

#### 3.2.3 Identification of Regions of Interest (ROIs)

For RSA and MVPA analyses, the ROIs were determined using the data from the passive session. We calculated the group average of the movies and rest contrast with a p-value threshold of p<0.001 in order to include the three levels of the AON (uncorrected). When selecting the borders of the ROIs (pSTS, parietal, and premotor cortices), we used the AAL atlas (Rolls et al., 2020) specific brain regions to mask the results. Then, we selected activated voxels within the atlas region. The brain regions used for masking are Temporal_Sup for pSTS, the union of Parietal_Inf and Parietal_Sup for the parietal cortex, and Precentral for the premotor cortex. The saved voxels for the premotor regions were found within the expected premotor activation based on previous studies (Ferri et al., 2015). The coordinates and cluster numbers of the extracted ROIs can be found in Table 1 and Figure 6.

**Table 1.** Cluster number, peak thresholds, and peak coordinates of the defined ROIs.

| Region of Interest | Left Hemisphere |  |  | Right Hemisphere |  |  |
| --- | --- | --- | --- | --- | --- | --- |
|  | Cluster Number | Peak Threshold | Peak Coordinates | Cluster Number | Peak Threshold | Peak Coordinates |
| pSTS | k=27 | t=4.98 | -54 -46 19 | k=114 | t=6.47 | 54 -42 19 |
| parietal | k=440 | t=9.11 | -28 -58 59 | k=459 | t=8.19 | 36 -54 56 |
| premotor | k=68 | t=5.95 | -36 -4 52 | k=17<br>k=22 | t=5.14<br>t=4.92 | 38 -4 49<br>50 2 46 |

**Figure 6.**
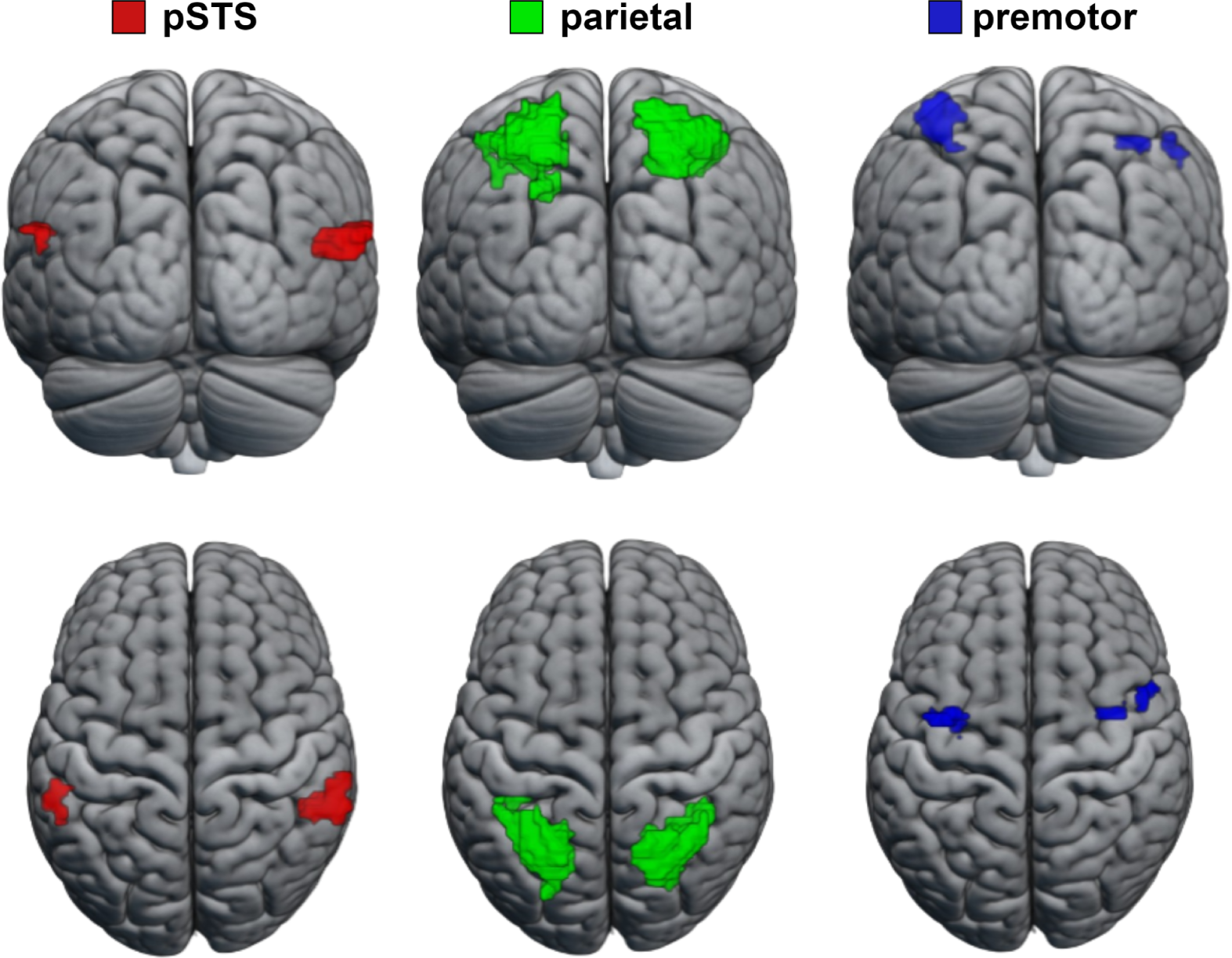
ROIs based on the group analysis.

### 3.3 RSA Results

Neural RDMs were generated by using correlation distance measure between each variable’s spmT for each ROI by averaging data from all subjects (Figure 7). Then, neural RDMs were compared with model RDMs. This comparison was carried out using a one-sided Wilcoxon signed-rank random-effects test corrected for multiple comparisons (FDR) at a threshold of p<0.05, employing Kendall’s τ as the correlation metric. The upper and lower bounds of the noise ceiling were determined by estimating a hypothetical model with the highest possible average correlation to the reference RDM, once again utilizing Kendall’s τ. Neural and model RDM comparisons are shown in Figure 8.

**Figure 7.**
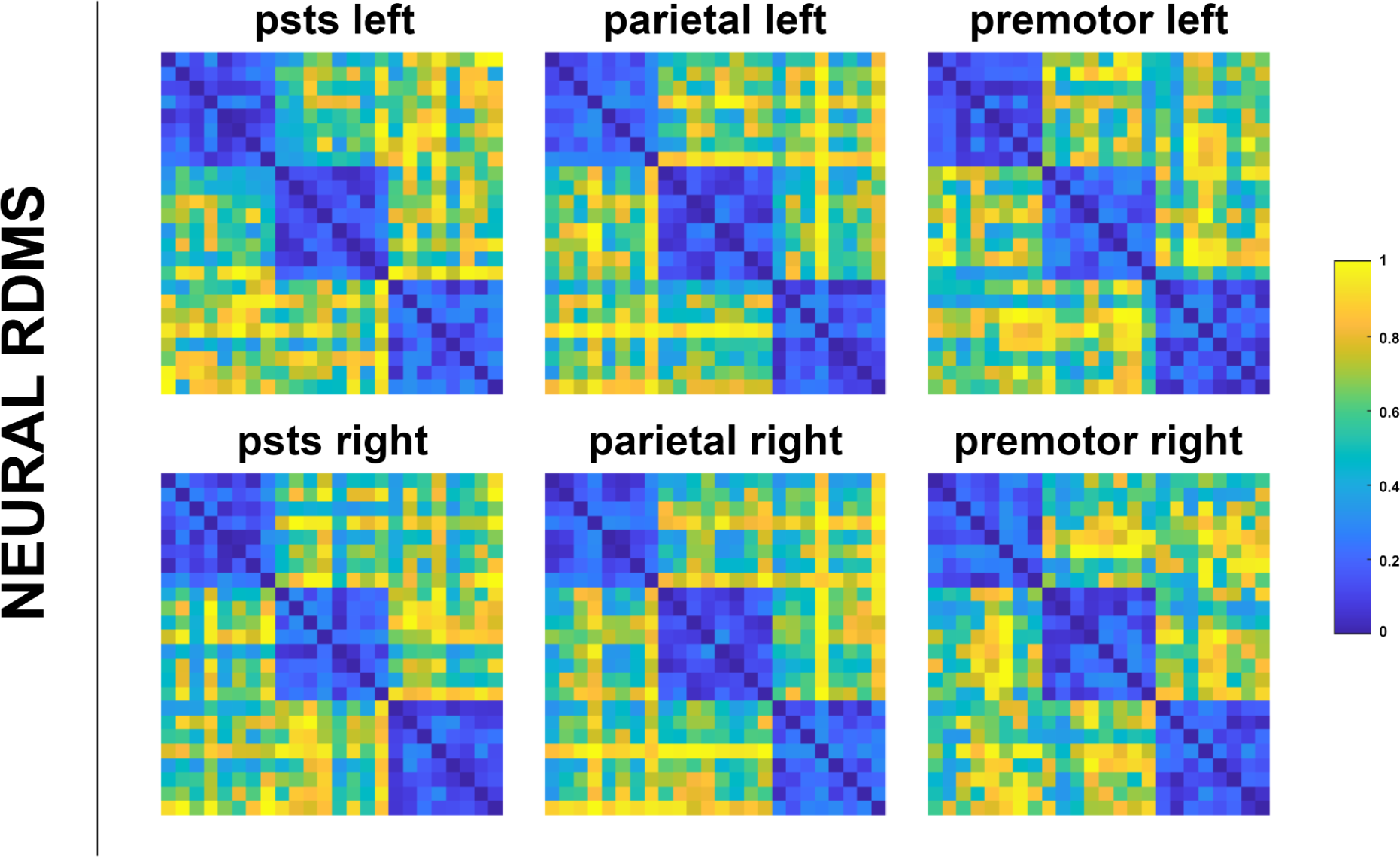
Neural RDMs averaged over all subjects. Each dissimilarity matrix (24 x 24) is separately rank transformed and scaled into [0,1].

**Figure 8.**
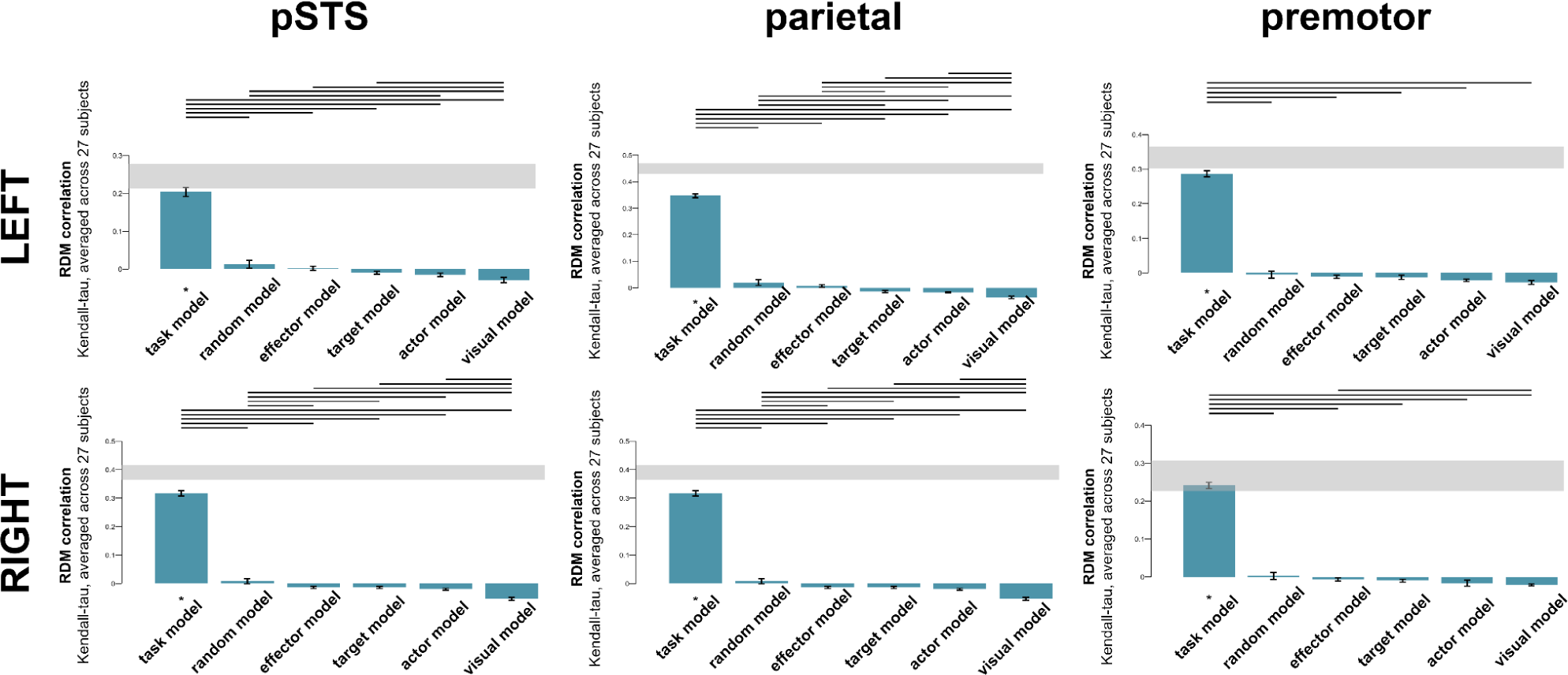
Comparison of neural RDMs and model RDMs. Gray bands over the graphs indicate the noise ceiling, horizontal black lines indicate significant differences between models, and (*) indicates significant similarity to the ROI RDM at p<0.05 FDR corrected level. Error bars indicate the standard error.

For all ROIs, only the task model was found to be significantly correlated with all neural RDMs (p<0.05, FDR corrected); none of the other models were significantly correlated with any ROI. Also, in the right premotor the task model’s Kendall’s τ value (0.2413) reached the noise ceiling (0.2266, 0.3068). Kendall’s τ and noise ceiling values for all ROIs and models can be found in Table 2.

**Table 2.** Statistical inference of comparison of neural RDMs and model RDMs. Kendall’s τ, Spearman’s ρ, and noise ceiling values are calculated.

| ROI | Actor Model |  | Effector Model |  | Target Model |  | Task Model |  | Visual Model |  | Random Model |  | Noise Ceiling |
| --- | --- | --- | --- | --- | --- | --- | --- | --- | --- | --- | --- | --- | --- |
| | Kendall's $\tau$ | p | Kendall's $\tau$ | p | Kendall's $\tau$ | p | Kendall's $\tau$ | p | Kendall's $\tau$ | p | Kendall's $\tau$ | p | |
| pSTS left | -0.0152 | 1.00 | 0.0020 | 0.43 | -0.0098 | 0.99 | **0.2038** | **0.00** | -0.0291 | 1.00 | 0.0126 | 0.08 | (0.2118,0.2784) |
| pSTS right | -0.0199 | 1.00 | -0.0133 | 1.00 | -0.0135 | 1.00 | **0.3171** | **0.00** | -0.0537 | 1.00 | 0.0084 | 0.11 | (0.3642,0.4157) |
| parietal left | -0.0174 | 1.00 | 0.0074 | 0.13 | -0.0138 | 1.00 | **0.3470** | **0.00** | -0.0362 | 1.00 | 0.0200 | 0.06 | (0.4289,0.4691) |
| parietal right | -0.0180 | 1.00 | 0.0031 | 0.66 | -0.0063 | 0.99 | **0.3122** | **0.00** | -0.0321 | 1.00 | 0.0094 | 0.13 | (0.3816, 0.4261) |
| premotor left | -0.0216 | 1.00 | -0.0113 | 1.00 | -0.0124 | 0.99 | **0.2870** | **0.00** | -0.0277 | 1.00 | -0.0049 | 0.69 | (0.3015,0.3654) |
| premotor right | -0.0211 | 1.00 | -0.0068 | 0.92 | -0.0101 | 0.99 | **0.2413** | **0.00** | -0.0165 | 0.98 | 0.0025 | 0.42 | **(0.2266,0.3068)** |

### 3.3 Decoding Analysis Results

“Female vs. male”, “hand vs. feet”, and “human vs. object” features were decoded under each task in each ROI, yielding a total of 54 different decoding analyses (2 hemispheres x 3 ROIs x 3 tasks x 3 features). Linear support vector machine classifiers were individually trained for each combination.

To assess decoding accuracy, we conducted one-sample t-tests comparing the decoding results from 27 subjects against the chance level (50%). Figure 9 displays accuracy minus chance results of all the decoding analyses conducted. Among the core regions of the AON, pSTS successfully decoded the target of the action (human vs object) in the target task on the left hemisphere and in the actor task on the right hemisphere. None of the other features were successfully decoded in pSTS under any task. On the other hand, the parietal cortex successfully decoded all features under various task conditions although there were some slight hemispheric differences. The actor of the action (female vs. male) was successfully decoded under the actor task in both hemispheres. The effector of the action (hand vs. feet) was successfully decoded in both hemispheres under all task conditions. The target of the action (human vs object) was successfully decoded in the target and actor tasks on the left hemisphere and in the target and effector tasks on the right hemisphere. In the premotor cortex, the actor of the action (female vs. male) was not decoded under any task condition. However, the effector of the action (hand vs. feet) was successfully decoded in the effector task in both hemispheres and in the actor task on the right hemisphere. In addition, the target of the action (human vs. object) was successfully decoded under the target task in both hemispheres.

**Figure 9.**
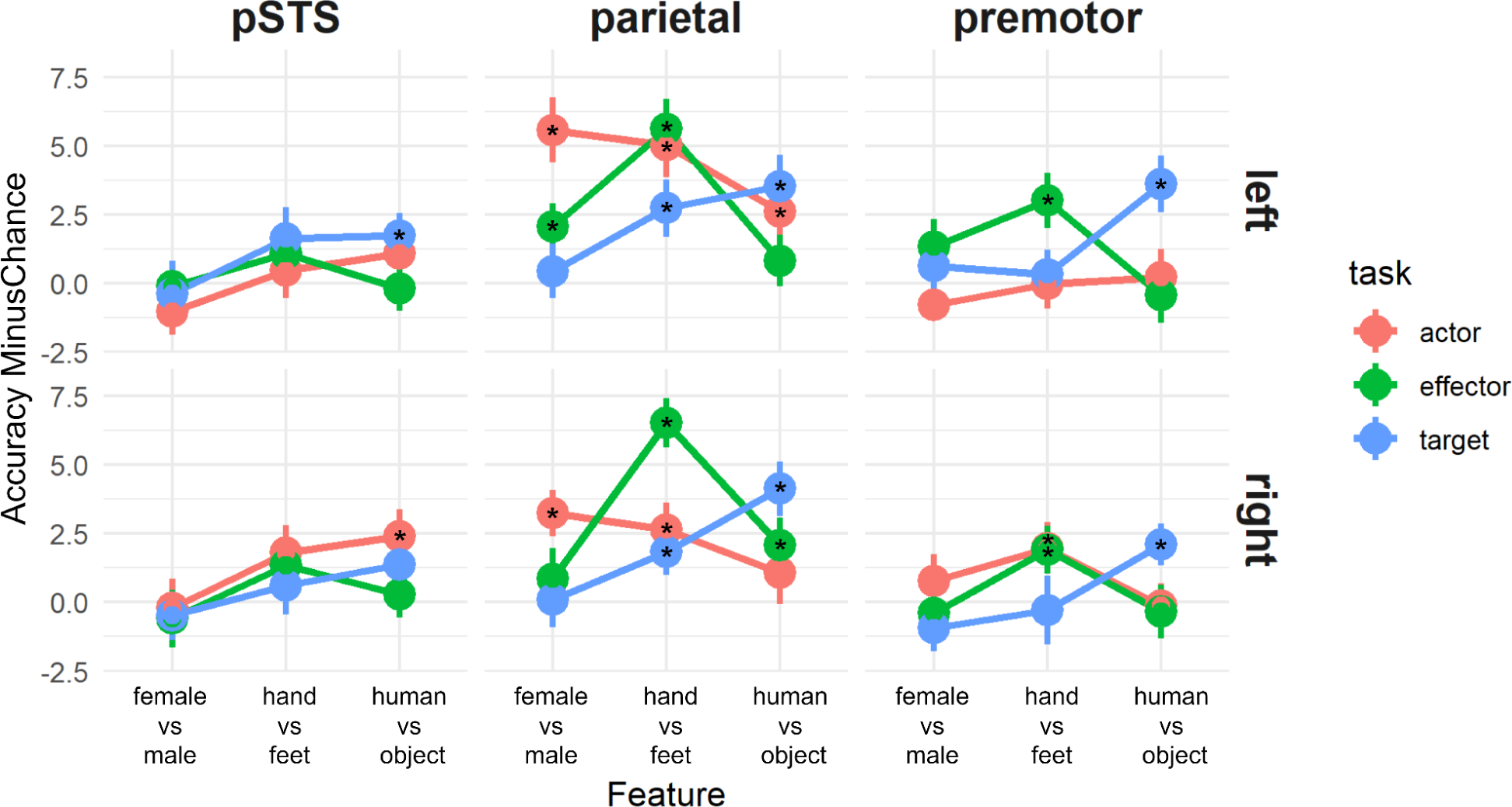
Accuracy minus chance results from the decoding analysis averaged over subjects. Significantly successful decodings are marked with “*”.

In order to examine the effect of factors on decoding accuracy and the differences between the conditions, we performed a 2 (Hemisphere) x 3 (ROI) x 3 (Task) x 3 (Feature) repeated measures ANOVA. The statistics for the main effects and interactions are summarized in Table 3.

**Table 3.** Results of the repeated measures ANOVA on decoding accuracy.

| 2 (Hemisphere) x 3 (ROI) x 3 (Task) x 3 (Features)<br>Repeated Measures ANOVA Results |  |  |  |
| --- | --- | --- | --- |
| Cases | F value | p value | $\eta^2$ |
| hemi | 1.433 | 0.242 | - |
| **roi** | **48.236** | **<0.001** | **0.038** |
| task | 0.130 | 0.879 | - |
| **feature** | **7.003** | **0.002** | **0.015** |
| hemi * roi | 0.905 | 0.411 | - |
| hemi * task | 0.526 | 0.594 | - |
| roi * task | 1.728 | 0.149 | - |
| hemi * feature | 0.581 | 0.563 | - |
| roi * feature | 1.535 | 0.197 | - |
| **task \* feature** | **6.702** | **<0.001** | **0.024** |
| **hemi \* roi \* task** | **3.443** | **0.011** | **0.007** |
| hemi * roi * feature | 0.332 | 0.856 | - |
| hemi * task * feature | 0.485 | 0.747 | - |
| **roi \* task \* feature** | **2.325** | **0.021** | **0.012** |
| hemi * roi * task * feature | 0.324 | 0.956 | - |

While there was no effect of hemisphere and task (See Table 3 for statistics), there was a main effect of feature (F = 7.003, p = 0.002, η²=0.015) and ROI (F = 48.236, p < 0.001, η²=0.038). Post-hoc tests showed that ‘hand vs feet’ decoding accuracy was significantly higher than ‘female vs male’ (MD=1.566, t=3.731, p=0.001 Bonferroni-corrected), with no other significant comparison results for the feature. For ROI, the parietal cortex’s decoding accuracy was significantly higher than the pSTS region (MD=2.227, t=8.684, p<0.001 Bonferroni-corrected) and premotor cortex (MD=2.133, t=8.316, p<0.001 Bonferroni-corrected). There was no significant decoding performance difference between the pSTS and premotor cortex.

There was also a significant interaction of task and feature (F=6.702, p < 0.001, η²=0.024). When attention was directed to a specific feature within a task, the decoding performance peaked across the related task (female vs. male in the actor task, hand vs. feet in the effector task, and human vs. object in the target task). Notably, within the effector task, decoding for hand vs feet surpassed both female vs. male (MD=2.727, t=3.956, p=0.004, Bonferroni-corrected) and human vs. object (MD=2.875, t=4.171, p=0.002, Bonferroni-corrected). Conversely, no significant distinction in feature decoding performances was observed within target and actor tasks.

Also, hemi*roi*task (F=3.443, p=0.011, η²=0.007) and roi*task*feature (F=2.325, p=0.021, η²=0.012) interactions were found significant. None of the other interactions were significant.

## 4. Discussion

In this study, we investigated how top-down attention influences the neural representations in the Action Observation Network, including the posterior superior temporal sulcus (pSTS), parietal cortex, and premotor cortex. Utilizing both univariate and multivariate analysis techniques, we examined how different tasks, attention to different features of an action, influence the representation of perceived actions within these ROIs. Our findings showed that tasks affect neural activity across all ROIs. Importantly, we observed that each task exerts a unique and distinctive influence on the activity within each specific ROI.

Univariate results showed that for both passive and active sessions, the AON was active. However, in the active session, we observe an extended neural activity, especially in the parietal cortex. Also, the premotor area remains inactive during the passive session, in contrast to its pronounced activation in the active session. The introduction of attentional demand during the observation of action videos significantly heightened spatial activation across AON. This finding is consistent with the results of Orban et al. (2019), who reported extended activation in the parietal and premotor cortex where participants engaged in tasks related to action features.

We also explored differences in brain activation between tasks. Each task’s contrast with the rest yielded very similar activity. Furthermore, we found no activation when contrasting tasks with each other (Actor-Effector, Target-Actor, etc.). Additionally, reaction times and accuracy rates showed no significant variations between tasks. To better understand the complex neural dynamics, we utilized multivariate pattern analysis alongside univariate analyses. Multivariate approaches have demonstrated higher sensitivity for detecting subtle patterns in neural activity (Haynes, 2015).

RSA results revealed that all ROIs in the AON show neural patterns that align with the ongoing task. The task model, but not the feature models, demonstrated significant similarity between neural RDMs for all ROIs. It shows that despite their distinct functions, the ROIs exhibit synchronization with the task, highlighting a shared adaptability to attentional demands.

Additionally, the right premotor region reached the noise ceiling, revealing that the task model alone is sufficient to account for neural patterns in this area. In contrast, none of the models reached the noise ceiling for the other regions. This may be due to the premotor region’s higher hierarchical position within the AON (Nelissen et al., 2011), indicating that tasks have a more significant impact on this region compared to the pSTS and parietal regions. While RSA results emphasize the task’s role in shaping AON activity, it’s important to acknowledge that distinct ROIs within the AON may inherently contribute to processing different action features (Urgen et al., 2019). To explore this, we further examined how well we could decode specific features within each task across these ROIs.

Decoding analysis revealed more detailed results regarding the effects of attention on the ROIs. First, we conducted ANOVA on the accuracy minus chance values. This analysis revealed no significant main effect on the task, indicating consistent cognitive demand and difficulty levels across tasks. Also, there is no main effect on the hemisphere, which shows that hemispheres exhibit no difference in terms of action observation processing. On the other hand, a main effect was observed for both ROI and feature, suggesting some levels of the AON may contain more information about the overall action features, and some features may be encoded more distinctly than others. The significant hemi * roi * task interaction suggests diverse impacts of tasks on ROIs across hemispheres. This interaction implies that ROIs in different hemispheres may respond differently to varying tasks, potentially influencing neural processing distinctively. Similarly, the observed effect of roi * task * feature implies differing task-induced effects on ROIs, potentially influencing the decoding performance of features. These interactions highlight the subtle and differential responses of brain regions to task variations, hinting at potentially distinct neural processing strategies based on the specific cognitive demands posed by different tasks.

Decoding results showed distinct patterns of neural activity within each ROI, indicating the specialized roles of different brain areas in action processing and extending previous work (Stehr et al, 2021; Shahdloo et al, 2022; Orban et al 2019). In the pSTS region, “female vs. male” and “hand vs. feet” classifications were unsuccessful across all tasks. However, it successfully decoded “human vs. object” in both actor and target tasks. This distinction is noteworthy because the object condition involves the actor manipulating an object, while the human condition involves the actor pushing another human, introducing a social dimension. Notably, the pSTS region plays a significant role in social processing, and this region is recognized as a social hub (Dasgupta et al., 2017). The distinction in social context between these conditions could be a factor contributing to the distinct neural presentations of the object and human videos within the pSTS region, potentially explaining why decoding was successful only for the “human vs. object” classification.

Decoding in the parietal cortex revealed an interesting result that decoding the “hand vs. feet” condition under all three tasks and for both hemispheres was successful. Several studies have shown that the parietal region is involved in encoding the effector. For instance, Errante et al., (2021) conducted research indicating that the parietal region can decode both grip type and the interaction between grip type and goal. This finding aligns with our successful decoding of “hand vs. feet” classification. The decoding accuracies across all features were significantly higher in the parietal region compared to other regions. The parietal cortex is known to encode the abstract meaning of actions, contributing to our understanding of the conceptual aspects associated with various movements (Gottlieb, 2007). Additionally, it plays a distinguished role in action discrimination, differentiating between different types of actions (Orban et al., 2019). Given that this region encodes abstract semantic information and demonstrates capability in discriminating actions, it is plausible that it requires the encoding of substantial information related to overall action features. This could potentially account for the enhanced decoding performance of the parietal cortex, enabling it to decode all features more effectively than other ROIs.

In premotor ROI, we observed that for the left hemisphere, “hand vs. feet” was successfully decoded under the effector task, and “human vs. object” decoding was successful under the target task. For the right hemisphere, “hand vs feet” is being successfully decoded under effector and actor tasks, and “human vs. object” is decoded under the target task. This suggests a pronounced task and feature interaction within this ROI. The premotor region appears to exhibit a greater sensitivity to tasks when compared with pSTS and the parietal cortex. This inclination may be attributed to the premotor region’s higher position in the AON hierarchy (Nelissen et al., 2011). Classification of “female vs. male” was unsuccessful across all tasks, indicating a lack of sensitivity to the actor in the premotor region. This can be explained by the fact that this region primarily encodes the kinematics of actions (Binkofski & Buccino, 2006; Takahashi et al., 2017), and variations in the actor do not affect the kinematic patterns. Also, in RSA analysis the only region that could reach the noise ceiling was the right premotor region, which also shows the premotor’s adaptation ability to the task at hand.

## 5. Conclusion

In this research we investigated the task based modulation of action perception on AON. We found that the type of the task affected the neural activity across all levels of the AON, with ROIs adapting their response patterns based on specific task requirements. Further analysis revealed task-specific effects on each ROI, interacting with action features. This reflects the hierarchical organization of ROIs, each residing at different levels and encoding distinct action features. Our findings highlight that top-down attentional modulation leads to alterations in the neural representation of action features within AON nodes. Importantly, these effects are observed uniquely across regions, each reflecting their initial encoding scheme and level of hierarchy in the AON network.

### Limitations

In the course of our investigation, certain limitations need to be acknowledged to provide a comprehensive understanding and potential implications of our study. Firstly, our study utilized a relatively small stimulus set consisting of 8 videos featuring a single action. While the observed activation patterns align with existing literature on action observation neural responses, the limited variety in stimuli may raise concerns about the generalizability of our findings to a broader range of actions.

Also, since the stimuli are quite complex due to our aim to reflect on real-life scenarios, the perfect control for low-level features was not possible. The subtle differences between the actor’s clothing and the transition between the starting and ending positions may vary even though it is minimal. However, we did not observe any similarities between the low-level visual model with the neural RDMs in RSA, which diminishes the possibility of any confounds.

Additionally, our study did not incorporate eye-tracking measures to assess participants’ gaze patterns during the task. The potential influence of eye movements, particularly in response to the dynamic nature of our stimuli, introduces a source of variability that we did not directly address (Schütz et al., 2011). Future research incorporating eye-tracking methodologies could shed light on the role of involuntary eye movements and their impact on neural activation patterns in similar experimental paradigms.

### Future directions

Building on our current findings, several promising paths for future research can be identified. A particularly promising direction involves enhancing the ecological validity of neuroscience research by incorporating more naturalistic stimuli such as real-world actors (Pekçetin et al, 2023). This transition towards real-world scenarios could provide a more comprehensive understanding of how action perception functions in complex, dynamic environments, beyond the constraints of simplified laboratory settings (Maselli et al., 2023).

Future research should prioritize a more in-depth exploration of how varying cognitive tasks distinctly shape neural responses within the domain of action perception. The selection of a specific task can profoundly influence neural activity, both globally and within specific brain regions. By systematically investigating the impact of different tasks, researchers can uncover task-specific neural signatures and gain deeper insights into the mechanisms underlying action perception. Additionally, a comprehensive understanding of the interplay between bottom-up sensory input and top-down cognitive modulation is crucial for developing sophisticated computational models of action perception.

Beyond the scope of cognitive neuroscience, a promising direction for future research lies in the development and real-world application of action perception algorithms. These algorithms, capable of recognizing and interpreting human actions, have significant potential to revolutionize various technologies. For instance, integrating such algorithms into drones could enhance their capabilities in search and rescue missions. This integration not only pushes the boundaries of our scientific understanding but also opens up exciting possibilities for improving human-technology interaction in critical and complex scenarios.

## Data Availability

The data is available upon reasonable request from the first author. The codes and the videos can be found in the GitHub profile of the first author: https://github.com/aslieroglu

## Acknowledgments

The authors wish to express their deepest gratitude to Aysu Nur Koç for help in data collection and manuscript review, to Murat Batu Tunca, and Ömer Rençbereli for their support in data collection and to Ali Feza Karahaliloğlu for manuscript review.

